# Double dissociation of spontaneous alpha-band activity and pupil-linked arousal on additive and multiplicative perceptual gain

**DOI:** 10.1101/2023.09.13.557488

**Authors:** April Pilipenko, Jason Samaha

## Abstract

Perception is a probabilistic process dependent on external stimulus properties and one’s internal state. However, which internal states influence perception and via what mechanisms remain debated. We studied how spontaneous alpha-band activity (8-12 Hz) and pupil fluctuations impact visual detection and confidence across stimulus contrast levels (i.e., the contrast response function or CRF). We found that weak pre-stimulus alpha power induced an “additive” shift in the CRF, whereby stimuli were reported present more frequently at all contrast levels, including contrast of zero (i.e., false alarms). Conversely, pre-stimulus pupil size had a “multiplicative” effect on detection such that stimuli occurring during large pupil states (putatively corresponding to higher arousal) were perceived more frequently as contrast increased. Signal detection modeling reveals that alpha power changes detection criteria equally across the CRF but not detection sensitivity (d’) whereas pupil-linked arousal modulated sensitivity, particularly for higher contrasts. Interestingly, pupil size and alpha power were positively correlated, meaning that some of the effect of alpha on detection may be mediated by pupil fluctuations. However, pupil-independent alpha still induced an additive shift in the CRF corresponding to a criterion effect. Our data imply that weak alpha boosts detection and confidence by an additive factor, rather than by a multiplicative scaling of contrast responses, a profile which captures the effect of pupil-linked arousal. We suggest that alpha-power and arousal fluctuations have dissociable effects on behavior. Alpha reflects the baseline level of visual excitability, which can vary independent of arousal.

**Significance statement:** Nearly a century ago, brain waves around 8-13 Hz (the “alpha-band”) were discovered and linked to visual processing and cortical arousal. However, the precise way that alpha activity shapes perception and relates to arousal is unsettled. We recorded pupillometry and EEG while subjects detected and reported confidence for visual stimuli with varying intensity. Stimuli occurring during states of strong alpha were seen less often, regardless of intensity level, suggesting alpha exerts subtractive inhibition on perception and confidence. Pupil size (a proxy for arousal) was found to correlate with alpha yet, surprisingly, has a different effect on perception. Small pupil lowered perceptual sensitivity more as stimulus intensity increased. Our findings reveal distinct effects of alpha activity and arousal on visual perception.

## 1. Introduction

It has long been recognized that human perception is shaped by internal states of an observer in addition to external stimulus properties (1). Cases of near-threshold perception, whereby the same weak physical stimulus is sometimes perceived and sometimes not, provide a compelling example. By studying the factors that predict trial-to-trial variability in the perception of a constant visual stimulus, much has been learned about which internal states contribute to shaping perception. Two decades of recent work have revealed that ongoing neural oscillations in the alpha-band (around 8-13 Hz) reliably predicts visual perception across a variety of studies and stimulus types (recently reviewed in (2)). Specifically, states of low ongoing alpha power have frequently been found to lead to an increased probability of stimulus detection and enhanced subjective judgments of awareness (3–18). However, the mechanisms by which ongoing alpha oscillations implement inhibition and impact perception remains debated.

In recent years, researchers have begun including stimulus-absent trials in addition to near-threshold stimulus present trials in experiments in order to estimate false alarm rates and compute the signal-detection theory (SDT) measures d’ and criterion (reflecting sensitivity to the stimulus and response bias, respectively). Most of these experiments have consistently found that low alpha power leads to a liberal shift in criterion, making observers more likely to report stimulus presence regardless of the actual presence of the stimulus, and have failed to find an effect of alpha on d’ (18–21).

Complementary findings come from numerous recent experiments using discrimination tasks paired with subjective ratings (such as visibility or confidence judgments), which find that low pre-stimulus alpha power leads to higher visibility and confidence ratings without corresponding changes in discrimination accuracy (17, 22–26). Collectively, these findings have motivated an understanding of alpha oscillations as reflecting the overall state of excitation/inhibition in sensory areas, irrespective of whether the neural population represents signal/correct or noise/incorrect stimulus features (27) -a proposal termed the baseline sensory excitability model (or BSEM; (2)).

A few recent findings are inconsistent with the above evidence, however. One study found that pre-stimulus alpha in visual areas (source-localized on the basis of a target-, but not noise-, related localizer) modulated detection d’. However, this effect was only found on the subset of trials in which the subject adopted a conservative criterion (28). Another study found evidence for both a d’ and criterion shift with changing alpha levels. The authors reported that spontaneously lateralized alpha biases the apparent contrast of contralateral stimuli while at the same time non-lateralized alpha alters contrast discrimination sensitivity (29).

Despite intensive research on the topic, these competing accounts (whether alpha impacts criterion or sensitivity) have rarely been studied across the range of stimulus intensity levels that define an individual’s psychometric function. As a result, the way in which ongoing alpha interacts with stimulus strength is unclear, yet understanding this could provide crucial insights into the mechanisms by which alpha impacts perceptual processing and ultimately manifests as either criterion or sensitivity changes. In a SDT framework, the function linking changes in stimulus contrast to stimulus detection (contrast response function or CRF), could undergo different types of alpha-related gain modulation (see Figure 1). An additive gain mechanism would add (when alpha is weak) a constant amount to the internal sensory distributions produced by different contrast levels (included zero contrast, or stimulus absent response distributions). As shown in the simulation in Figure 1A, this would manifest as a parallel leftward shift in the CRF, a similar parallel shift in confidence judgments, a downward (or liberal) shift in criteria which is constant across contrast levels, and no change in detection sensitivity (Figure 1A). Alternatively, for alpha to impact sensitivity, a selective boost in signal over noise would need to occur, which can be captured in the multiplicative gain model. According to multiplicative gain, the effect of weak alpha would be to boost response distributions in proportion to their input strength such that weak responses (like when stimuli are absent or sub-threshold) would receive a negligible boost and stronger (higher contrast) stimuli would receive larger and larger boosts. Behaviorally, this would manifest as an increase in stimulus detections for higher contrast stimuli when alpha is low, a selective boost in confidence for high contrast stimuli, and a change in sensitivity that grows with stimulus contrast (Figure 1B). To date, only one study attempted to adjudicate these models, however they did not include stimulus absent trials, precluding a SDT analysis (30).

**Figure 1.**
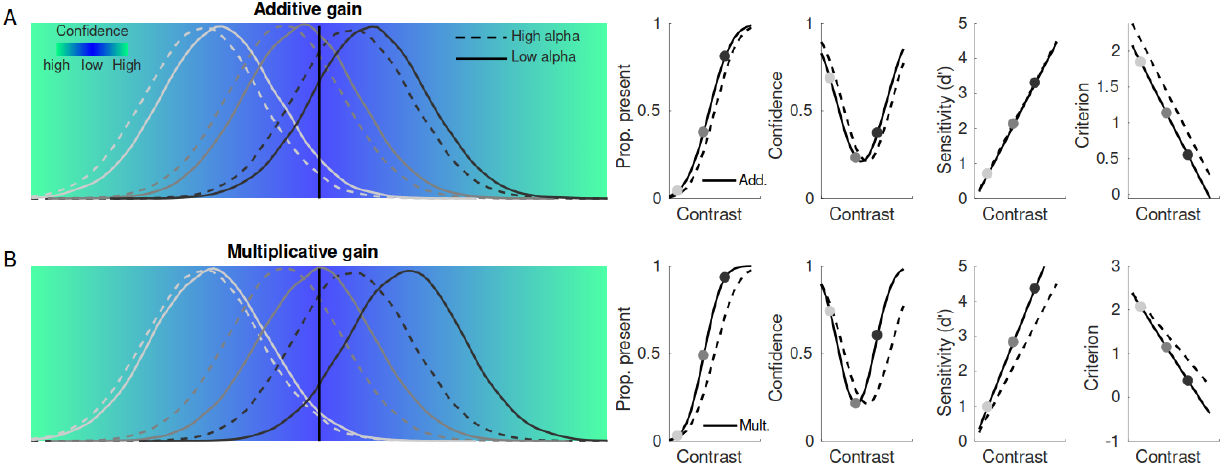
SDT interpretation of the psychometric function and confidence. Each stimulus intensity level gives rise to an internal response which can be characterized as coming from a Gaussian distribution. When an internal response on a given trial surpases the decision criterion (vertical line in panels A and B), the observer reports that a stimulus was present. Here, three stimulus intensity levels are highlighted as three distributions with increasing means (from light to dark lines). A standard framework for confidence in SDT is to assume that confidence on a given trial is a function of the “distance-to-criterion” of a sample of evidence, illustrated here in the color gradient. We simulated detection reports, confidence, sensitivity and criterion across varying stimulus intensities for two different states of pre-stimulus alpha power (the solid line representing low alpha power). **A)** Behavioral predictions under an additive gain model whereby low alpha adds the same constant to each distribution regardless of stimulus intensity. **B)** Predicted behavior under multiplicative gain, whereby the mean of each distribution is multiplied by a constant, producing larger increases as stimulus intensity grows.

**Figure 2.**
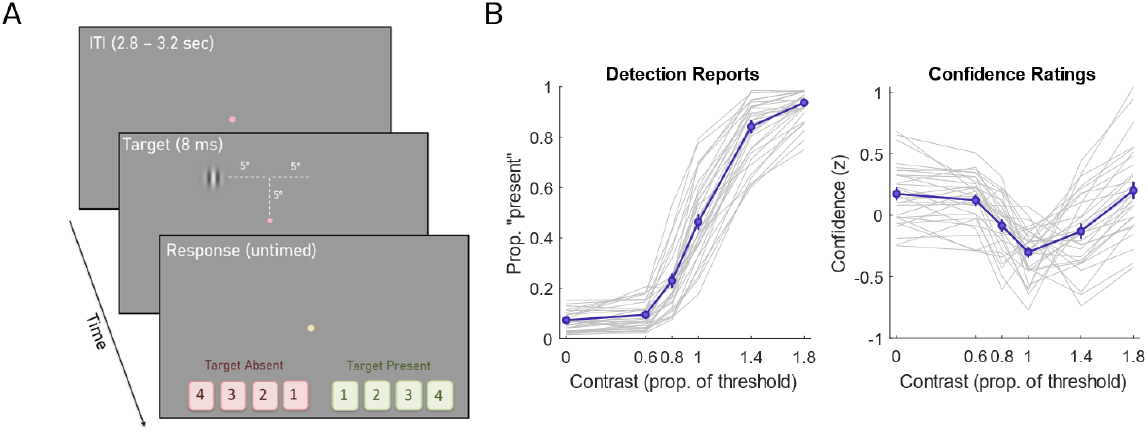
Task schematic and behavior. **A)** After a variable inter-trial interval (ITI) a brief Gabor patch (“target”) was presented either in the upper left or upper right visual field and observers (n = 30) reported whether they detected it or not along with their confidence, using a single button press (note that the dashed lines and text in this schematic were not displayed to observers). The target varied in contrast randomly from trial-to-trial between 0 (absent) and 1.8 times each individual’s 50% detection threshold (estimated prior to the main task via an adaptive procedure). **B)** The left panel shows the contrast response function (CRF) linking stimulus contrast to the proportion of “present” reports for each observer (gray line) and the group average (purple line). Right panel shows the mean confidence rating (z-scored) at each contrast level for individual observers and the group. The U-shaped confidence curve indicates that confidence increased as the stimulus became more clearly present or more clearly absent; a contrast of 1 indicates the 50% threshold level where uncertainty was highest and confidence was appropriately rated as lowest. Note that in all plots we display normalized confidence to more clearly see within-subject variation but all statistical tests were performed on raw confidence ratings. Error bars are ±1 SEM.

A final important factor, neglected by most prior work, is the potential influence of other perceptually-relevant factors that may co-vary with ongoing alpha power. Specifically, recent work has pointed to a positive correlation between spontaneous changes in alpha power and contemporaneous changes in pupil size, putatively reflecting arousal-linked brainstem neuromodulatory systems (31–35). Pupil changes correlated to ongoing alpha could influence perception in several ways, and thus could confound interpretation of an alpha effect. First, pupil increases are generally taken to reflect increases in arousal, which may enhance performance (36–39), and in vision studies in particular, increasing pupil size provides more lightfall on the retina, which increases detection in peripheral vision, but may lead to image blur and worsened fine discrimination (40, 41). Thus, recent work has highlighted the need to investigate alpha fluctuations that are independent of contemporaneous pupil changes (34, 42).

In the present study, thirty observers were asked to detect and report confidence in stimuli that varied across a range of contrast levels tailored to each individual’s CRF while alpha activity and pupil-diameter were simultaneously monitored. Across four metrics of behavior (detection, confidence, sensitivity, and criterion), we found that pre-stimulus occipital alpha power exerted an effect most consistent with additive, rather than multiplicative gain. In direct contrast, pre-stimulus pupil fluctuations lead to multiplicative but not additive effects on behavior. Importantly, controlling for the influence of pupil size on detection, alpha was still found to exert an additive effect, consistent with criterion but not sensitivity changes.

## 2. Results

We first determined which frequencies in our data were relevant for predicting perception by sorting trials according to whether the stimulus was reported as “seen” or “unseen” (collapsing all contrast levels) and testing for differences between these two states across the pre-stimulus power spectrum (from 3 – 40 Hz) derived from posterior channels with maximal alpha power (see Methods). As seen in Figure 3A, stimuli which were reported as “seen” showed significantly lower power in a relatively narrow frequency range from 6-14 Hz as compared to “unseen” stimuli. The fact that this effect was narrow-band in nature supports the claim that oscillatory alpha activity predicts stimulus detection. To further interrogate the mechanisms by which alpha influences detection across the CRF, we estimated pre-stimulus alpha power on single trials and computed four behavioral metrics: the proportion of “seen” responses, mean confidence ratings, sensitivity (d’), and criterion (c) for high and low pre-stimulus alpha power bins and for each contrast level.

**Figure 3.**
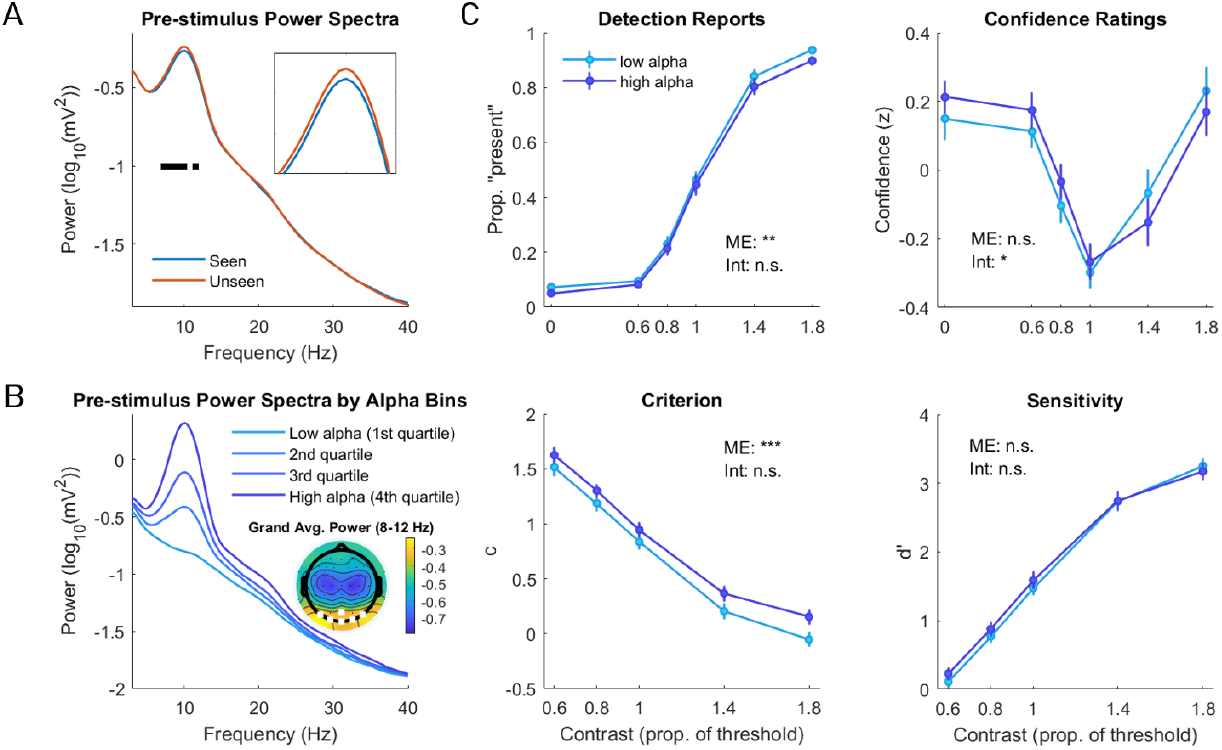
Additive effects or pre-stimulus power on behavior. **A)** The grand average pre-stimulus power spectrum from occipital channels reveals higher alpha power preceding unseen compared to seen trials, collapsing across all contrast levels. Black squares represent statistically significant differences (FDR-corrected), showing a relatively narrow-band effect limited to the alpha range (magnified in the inset figure). **B)** The grand average power spectrum for each alpha bin used in our analyses. The inset topoplot displays the grand average 8-12 Hz power with white dots indicating the electrodes used for all subsequent analyses. **C)** Detection reports showing the proportion of trials reported “present” by contrast level and pre-stimulus alpha power bin. Confidence depicts the average (z-scored) confidence rating of each contrast level for both high and low pre-stimulus alpha states. Criterion and sensitivity (d’) are shown in the two lower graphs. Overall, states of low pre-stimulus alpha power induced a main effect (ME) on detection reports without an interaction (int), boosting detection approximately equally across contrast levels leading to a constant criterion shift at all contrast levels and no interaction or impact on d’. Weak pre-stimulus alpha also translated the confidence curve leftward, boosting confidence for supra-threshold stimuli and reducing it for sub-threshold contrasts, thus leading to a significant interaction between alpha and contrast. The effects of ongoing alpha on all four behavioral markers best resemble those predicted by the additive model. Error bars are ±1 SEM. *denotes p<0.05, **denotes p<0.01, ***denotes p<0.001.

### 2.1 Pre-stimulus alpha power

We first analyzed detection reports (i.e., the proportion of “seen” responses) as a function of alpha power (high, low) and stimulus contrast (6 levels) using a 2-by-6 repeated-measures ANOVA. The additive model predicts that weak pre-stimulus alpha increases detection across all contrast levels equally, corresponding to a main effect in an ANOVA framework without an interaction. In contrast, the multiplicative model predicts that alpha power interacts with contrast such that the boost in detection during weak alpha is stronger for high compared to low contrasts.

As seen in Figure 3C, we observed a modest though approximately equal effect of alpha on stimulus detection across all contrast levels, leading to a significant main effect of alpha on the proportion of “seen” responses (F(1, 145) = 10.44, p = .003). The direction of this effect is consistent with literature showing an inhibitory function of alpha, whereby states of high pre-stimulus alpha power lead to lower detection reports (2). In contrast with the predictions of the multiplicative model, we found no evidence for an interaction between alpha power and contrast level (F(5, 145) = 0.53, p = .75), which suggests that pre-stimulus alpha power exerts an effect on detection that is uniform across contrast levels. Taken together, the influence of alpha on detection across the CRF is more consistent with an additive, rather than the multiplicative account of alpha.

The confidence rating data we collected can also help arbitrate between additive and multiplicative effects since, according to a standard SDT implementation of confidence (see Figure 1), additive gain should shift the U-shaped confidence curve rightward by a constant when alpha power is low. This would mean that, when alpha is strong, subjects feel more confident for weak stimuli (i.e., more confident that they are not detecting) and less confident for strong stimuli (i.e., less confident that they are detecting). This pattern should manifest as “cross-over” interaction in an ANOVA since the direction of the alpha effect flips depending on the contrast level. Multiplicative gain also predicts an interaction between alpha and contrast level although with a qualitatively different pattern whereby the effect of alpha is largely restricted to higher contrast levels (i.e., not a “cross-over” pattern).

As seen in Figure 3C, the effect of pre-stimulus alpha on confidence appears to flip in direction right around the threshold level of contrast such that participants are more confident in weak stimuli and less confident in strong stimuli when alpha is weak. This cross-over pattern resulted in a significant interaction effect (F(5, 145) = 2.71, p = .023), and a non-significant main effect of alpha (F(1, 145) = 0.59, p = .449). The pattern of the effect of alpha on confidence across the CRF most closely resembles that of the additive model.

Based on our simulation (Figure 1), the clearest dissociation between the additive and the multiplicative models is their predicted effect on sensitivity (d’) and criterion (c). Since the additive model equally boosts the HR and the FAR, no change in d’ is expected at any contrast level (no main effect or interaction), however, a change in criterion is expected and this effect should be approximately equal across contrast levels (a main effect, no interaction). On the other hand, the multiplicative model predicts an interactive effect of alpha on d’ and criterion, such that during states of weak alpha sensitivity is boosted more the higher the contrast. Our pattern of ANOVA results precisely follows that of the additive model predictions.

Specifically, our results showed a significant main effect of alpha power (F(1, 116) = 15.82 p < .001) on criterion, accompanied by no interaction effect (F(4, 116) = 1.49, p = .211), suggesting that the impact of alpha on criterion was approximately constant across the CRF. This effect can be seen in Figure 3C and the direction is consistent with prior work suggesting that weak alpha power induces a more liberal criterion, making participants report seeing stimuli more frequently, regardless of their actual presence.

Our analysis on sensitivity did not reveal a significant main effect of alpha power, F(1, 116) = 0.31, p = .583, nor interaction between alpha and contrast level, F(4, 116) = 1.49, p = .211. These results are visible in Figure 3B by the overlapping d’ measurement at each contrast level for both high and low pre-stimulus alpha power. Our results show that spontaneous states of high and low pre-stimulus alpha power did not influence sensitivity, even at higher contrast levels.

Taken together, these results provide additional support that spontaneous pre-stimulus alpha power changes detection rate by way of a criterion shift and not a sensitivity shift. Moreover, the effect of alpha power on criterion appears equal across the CRF, supporting the additive model.

In sum, the pattern of alpha effects across all four behavioral indicators were predicted best by the additive gain model. States of low alpha led to a uniform increase in detection reports inducing both HR and FAR increases, confidence ratings showed a cross-over interaction which depended both on alpha power and stimulus intensity, criterion showed a uniform change across stimulus intensities (with low alpha having a relatively more liberal criterion), and sensitivity was unaffected by changes in alpha at any contrast level.

### 2.2 Generalized linear model for pre-stimulus alpha power

The previous analysis considered the highest and lowest alpha power quartiles which somewhat arbitrarily classifies trials as “high” and “low” alpha trials while excluding all other trials. This approach aided in visualization of the effects, ease of interpretation, and consistency with prior literature (18, 19, 27), however we wanted to confirm that these effects were not due to the specific binning procedure used and still hold when utilizing the complete data set. We therefore implemented an alternative measurement of SDT using a generalized linear model (GLM) with a probit link function (43). Equivalent SDT measurements of c and d’ can be computed in a GLM framework (see *Materials and methods*), though they are somewhat less standard in the literature.

The results of the GLM analysis supported our initial findings and provided reason to believe that our binning procedure did not mask any potentially weaker effects nor create any artifactual ones. Parameter estimates corresponding to the effect of pre-stimulus alpha power on criterion and sensitivity were derived separately for each contrast level and participant. To assess the main effect of alpha power on c and d’ we collapsed over contrast levels and used a t-test to determine if the parameter estimates were different from zero. This revealed a statistically significant change in criterion (t(29) = 3.191, p = .003, 95% CI: [0.040, 0.187]), but not in d’ (t(29) = 0.546, p = .59, 95% CI: [-0.133, 0.231]). A one-way ANOVA assessed whether the parameter estimates capturing the effect of alpha on criterion and sensitivity changed with contrast level and found no statistically significant effects (criterion: F(4, 29) = 1.99, p = 0.1; d’: F(4, 29) = 2.01, p = 0.097), indicating a lack of an interaction and a relatively constant influence of alpha across the CRF.

### 2.3 Pupil size

We next sought to better understand how spontaneous pupil fluctuations may have affected perception. The pupil responds both to external stimuli (e.g., luminance changes) as well as internal states (e.g., relaxation) and is often used as a proxy for arousal, termed pupil-linked arousal (44, 45) As our analysis focused on pre-stimulus pupil states, just as in the case of alpha, single-trial variation in pupil size from our data likely reflects changes in internal cognitive states rather than external stimuli. This allowed us to address whether internally-driven states of pupil-linked arousal impacted stimulus detection in our paradigm and via what mechanisms (additive or multiplicative gain).

To estimate single-trial pupil diameter, we averaged over the same time window of data (−450ms to 50ms relative to stimulus onset) and implemented the same binning procedure used for the alpha analysis, which classified the pupil size high and low pupil states based on the first and fourth quartiles. We then analyzed all four behavioral measures (detection, confidence, criterion, and sensitivity) using repeated-measures ANOVAs with pupil size (high and low) and contrast level as predictors.

Figure 4A shows the effect of pre-stimulus pupil size across all behavioral metrics. In contrast to the effect of alpha, we observed a significant main effect of pupil size on detection (F(1, 145) = 16.3, p < .001) as well as a significant interaction with contrast level (F(5, 145) = 8.32, p < .001). This effect shows that on trials when participants happened to have a larger pupil size, they reported perceiving the stimulus more often but only for higher contrast levels (i.e., participants were not producing more false alarms).

**Figure 4.**
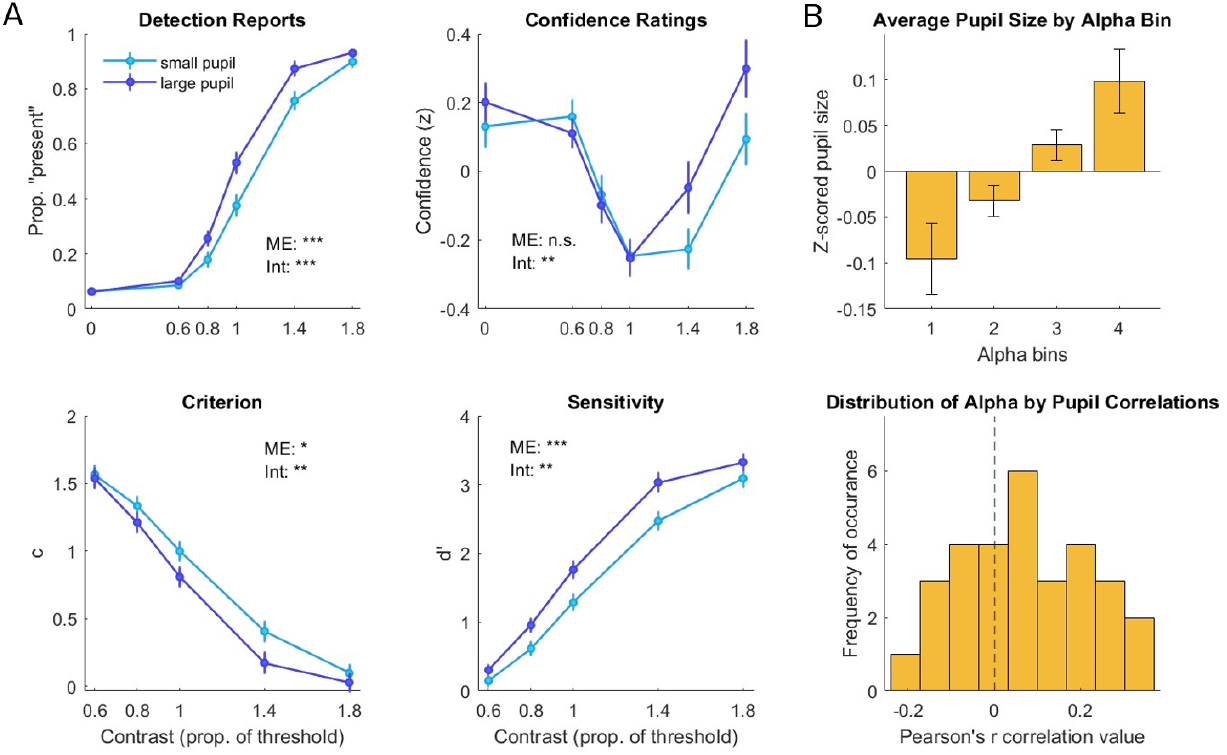
Multiplicative effects of pre-stimulus pupil size on behavior **A)** Large pre-stimulus pupil lead to more frequent stimulus detection, particularly at higher contrasts, without increasing false alarms. This effect corresponds to a boost in sensitivity that interacts with contrast level and is mirrored in a criterion shift with similar profile. Confidence on high pre-stimulus pupil trials was also selectively boosted at supra-, but not sub-threshold contrasts, as predicted by multiplicative gain. **B)** The top panel shows the group-level, z-scored pre-stimulus pupil size for each pre-stimulus alpha power quartile, depicting a positive linear relationship between spontaneous alpha and pupil size. The lower panel depicts the distribution of single-trial correlations between pre-stimulus alpha and pupil for each subject, which tends towards positive correlations. Error bars are ±1 SEM. *denotes p<0.05, **denotes p<0.01, ***denotes p<0.001.

This pattern of results is more consistent with pupil exerting a multiplicative as opposed to additive effect on the CRF. Profiles of the other behavioral metrics bore this out as well.

Specifically, large pupil lead to a boost in confidence ratings but primarily for stronger stimulus intensities, resulting in a significant interaction effect (F(5, 145) = 3.28, p = .008) and no main effect (F(1, 145) = 2.92, p = .1). Large pupil also led to an apparent liberal shift in criterion which was more pronounced for higher contrast levels, leading to a significant interaction effect (F(4, 116) = 3.72, p = .007) as well as a main effect (F(1, 116) = 5.77, p = .023). This criterion effect, however, could be understood to originate from a multiplicative change in sensitivity since, as the distributions increase along the internal evidence axis they become further from the criterion (producing an apparent shift in the “relative” criterion measured by standard SDT; see Figure 1). This seems to be the case as an interactive effect of pupil on d’ was clear, whereby large pupil increased sensitivity particularly at high contrasts (F(4, 116) = 3.72, p = .007). A significant main effect of pupil on d’ was also evident (F(1, 116) = 17.12, p < .001). In other words this apparent shift in criterion could simply be the result of highly similar FAR between the two pupil states with differing HR as is predicted by the multiplicative gain model in Figure 1B. Thus, whereas spontaneous changes in alpha power produce an additive effect on the CRF that alters criterion, pupil changes occurring at the same moment in time lead to a multiplicative effect that boosts sensitivity.

We next sought to understand how these opposing effects might be related. This question has particularly interesting theoretical implications since both spontaneous pupil fluctuations as well as spontaneous alpha power are often taken as proxies of internal states of arousal or attention, respectively, that are predictive of task performance(34, 35, 37, 39, 46, 47). To this end, we used the same alpha quartiles as used in the previous analysis of behavior but instead computed the normalized pre-stimulus pupil size at each alpha level. As seen in Figure 4B there is an apparent positive linear relation between spontaneous alpha fluctuations and contemporaneous pupil changes, with a significant increase in pupil at the highest compared to the lowest alpha level (t(29) = -2.759, p = .01, 95% CI: [-0.337, -0.050]). This relationship was corroborated by a single-trial correlation analysis whereby alpha and pupil were correlated on single trials separately for each participant. As shown in Figure 4B, most subjects had a positive correlation, the distribution of which was significantly different from zero at the group level (t(29) = 2.861, p = .008, 95% CI [0.022, 0.133]).

The strengths of the single-trial correlations were quite small, yet the direction of this effect is somewhat surprising. Namely, states of higher alpha power are generally related to states of larger pupil size, which suggests that these two positively correlated indices of internal states actually have opposing effects on behavior, given the previously observed results. That is, while states of low alpha power have higher detection on average and an additive scaling of the CRF, states of low pupil have lower detection on average and a multiplicative scaling of the CRF, yet the two are positively related. We therefore reasoned that the correlated and contemporaneous pupil effects could perhaps be influencing the observed effect of alpha on detection behavior, which is not something most other studies have controlled for.

### 2.4 Residual alpha power

The goal of this analysis was to see whether or not the effects of alpha power on detection was modified by alpha’s relationship with pupil size. Residual alpha power was estimated by taking the single-trial residuals of a linear model predicting pre-stimulus alpha power using pre-stimulus pupil size. The residuals of this model were then binned in the same manner as alpha power, i.e., 4 bins of ascending power and the analysis similarly considered the two most extreme bins (high and low) as predictors of behavior.

Consistent with our initial findings, we observed a significant main effect of residual alpha power on detection (F(1, 145) = 10.78, p = .003) with no evidence of an interaction (F(5, 145) = 0.38, p = .861; Figure 5B). This suggests that, in our data set, measurements that better isolate alpha power, such as controlling for pupil-linked arousal, still provide evidence for an additive effect of alpha on perception. Moreover, the cross-over effect of alpha on confidence ratings held when residualizing for pupil (see Figure 5B), showing a significant interaction effect (F(5, 145) = 2.7, p = .023) and no main effect (F(1, 145) = 0.41, p = .527). As further predicted by the additive model, residual alpha had a main effect on criterion (F(1, 116) = 19.97, p < .001) coupled with a non-significant interaction (F(5, 116) = 1.05, p = .387) and no main effect (F(1, 116) = 0.29, p = .591) or interaction (F(5, 116) = 1.05, p = .387) on d’.

**Figure 5.**
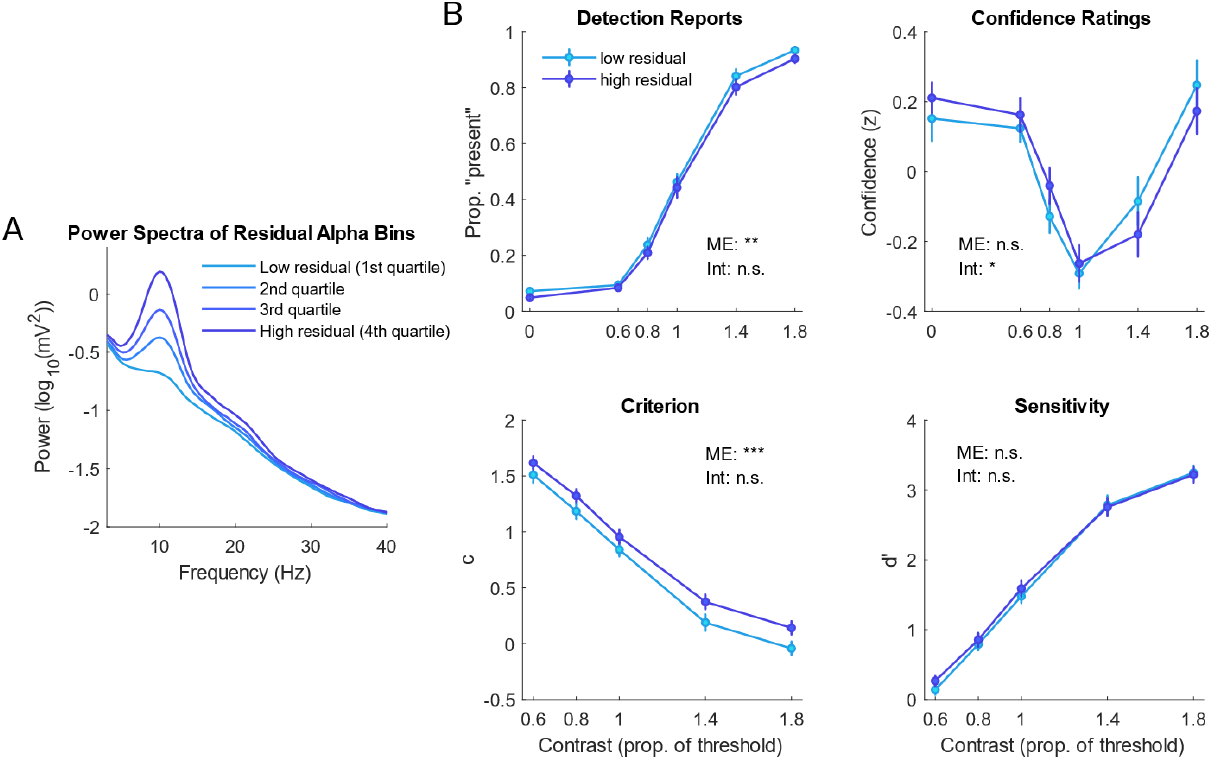
Pupil-independent pre-stimulus alpha fluctuations exert additive gain. **A)** The grand average pre-stimulus power spectrum for trials in each pupil-independant alpha quartile. **B)** The effects of residual (pupil-independent) alpha power on detection, confidence, criterion, and sensitivity are indicative of additive gain with a main effect on detection reports and criterion, a contrast-by-alpha interaction on confidence, and no effect of alpha on d’. Error bars are ±1 SEM. *denotes p<0.05, **denotes p<0.01, ***denotes p<0.001.

## 3. Discussion

Gain modulation of sensory responses allows the brain to dynamically shape how it responds to external stimuli as a function of ongoing internal brain states. Alpha-band oscillations are the most prominent form of intrinsically generated brain activity seen in the human EEG and, despite having been discovered nearly a century ago (48), the precise modulatory effect of alpha activity on perception has remained unclear. Here we show that states of strong (weak) ongoing alpha activity suppress visual detection via a constant subtractive (additive) factor, corresponding to a reduction of stimulus detection that is approximately equal across all stimulus intensity levels. In direct contrast, we find that pupil fluctuations, known to partly reflect arousal-related neuromodulatory activity, multiplicatively scale visual detection, such that large pre-stimulus pupil states boost visual detection more for higher contrasts. Importantly, isolating fluctuations in alpha power from contemporaneous pupil changes still led to the observation of an additive effect of low alpha on detection.

Our findings have important implications for understanding how ongoing alpha activity shapes perceptual behavior. The additive gain we observed suggests that states of low alpha should not lead to any change in perceptual sensitivity (d’) since low alpha boosts detection reports even in the absence of any stimulus (i.e., false alarms). This is indeed what our SDT analyses revealed and is congruent with multiple recent studies showing an effect of alpha on detection criterion, but not sensitivity (18–21, 49). Our results build on this prior literature by showing that the effect of alpha on detection criterion is not restricted to just the case of threshold stimuli but is also observed across the range of contrast levels spanning each individuals CRF, and also by showing that the criterion effect holds after controlling for the correlation between ongoing alpha and pupil size. Our findings also shed light on the computations underlying confidence in visual detection. According to some accounts, confidence is computed as the probability of being correct, which should increase along with d’ (50–53). However, our results show that confidence increases along with alpha-induced criterion shifts even with no accompanying change in d’. This could naturally come about if observers use a “distance-to-criterion” computation as illustrated in Figure 1. When alpha is low and an additive factor is applied equally to stimulus present and absent distributions, high contrast stimuli become further from the decision criterion (leading to higher confidence) and low contrast/absent stimuli get closer to the criterion (leading to lower confidence). Thus, our analysis of pre-stimulus brain states adds to the growing body of behavioral evidence that confidence is not computed as the Bayesian probability of being correct (51, 54–63).

Collectively, our findings support the idea that ongoing alpha power reflects the overall excitability of the visual cortex, a concept which is at least partly dissociable from the notion of arousal, since we find the two have differing effects on perceptual behavior. In line with the recently proposed BSEM (2), the alpha-induced change in cortical excitement can be described as a baseline shift, which is hypothesized to influence signal-detecting neural populations even in the absence of any stimulus. In this scenario, the base increase would push the firing rate closer to the detection criterion, increasing the chances of perceiving a weak stimulus (i.e., a hit), while simultaneously increasing the chances of weakly perceiving something which was not there (i.e., a false alarm). In contrast, we propose that the spontaneous changes in pupil size can be taken as a proxy of internal arousal, partly distinct from changes in visual cortical excitement. Pupil-linked arousal is widely believed to be modulated by the locus coeruleus-norepinephrine (LC-NE) system and/or the superior colliculus (SC; 36, 44, 45, 54), and may be relatively unlinked with stochastic variation in visual cortical excitability. The effect of pupil-linked arousal on behavior exhibited a gain response which could involve multiple cellular mechanisms including changes in synaptic input or membrane conductance (65). On the other hand, the additive impact of alpha on detection may be explained by different cellular mechanisms such as shunting inhibition (65).

The results of this study are consistent with many recent findings showing that spontaneous alpha power influences detection by way of a criterion shift and not sensitivity. Yet, a handful of studies have reported that spontaneous alpha power may increase sensitivity in certain cases. In addition to finding an effect of alpha on subjective contrast appearance, Balestrieri and Busch (2022) also observed that weak pre-stimulus alpha led to an increase in contrast sensitivity in spatial 2-alternative forced choice contrast discrimination task (29). One notable difference from the present study is that Balestrieri and Busch used a continuously running staircase to modify stimulus contrast throughout the main task. Although speculative, this methodological choice could have introduced complex trial-by-trial dependencies between states of fluctuating alertness and subsequent task difficulty, perhaps leading to spurious correlations between behavior and spontaneous brain states. In our study we ensured that our predefined stimulus levels were truly identical by fixing contrast levels at the beginning of the main task based on a pre-task staircase.

Zhou et al. (2021) similarly found an effect of alpha power on contrast sensitivity in a discrimination task querying whether a backward-masked stimulus had been a grating or noise (28). The effect on d’ was found in only one of their conditions when a conservative detection criterion was experimentally induced and did not replicate in their liberal criterion condition. Notably, their analysis of pre-stimulus power targeted brain areas which were feature-specific to the target stimulus, which differs from papers looking at global alpha changes (which we presume reflect target and non-target excitability fluctuations). Thus, it is possible that isolating feature-specific areas to assess alpha power might not capture overall visual cortical excitability and instead might capture the local variability of a target-related population. If this were the case, then it would follow that sensitivity shifts could arise from changes in feature-specific alpha power, as they do when alpha is topographically modulated by attention, for example (66–70).

Lastly, one other study investigating alpha power’s role in perception across the CRF presented results which are at odds with our findings. Chaumon and Busch (2014) showed evidence of a multiplicative effect pre-stimulus alpha power on detection (30). This study used a go/no-go task paradigm to report stimulus presence which means participants only responded when they perceived the stimulus (and withheld a response when they did not) and they did not include stimulus-absent trials and thus could not compute SDT measures. We suggest that task demands in their study may explain these inconsistent results. Since the go/no-go paradigm requires additional effort (albeit quite small) to actively report “seen” as opposed to doing nothing to passively report “unseen”. This may have created an additional bias such that participants were less inclined to report stimulus presence at low contrasts but when the stimulus was more obvious (at high contrasts) they were more willing to engage in the more effortful behavior, leading to a multiplicative-like pattern. Overall, the literature would benefit not only from additional studies measuring the perceptual effects of alpha across the CRF but also from studies investigating potential differences in the effects of alpha during specific paradigms or task demands.

Much about the stochastic processes underlying perception is still unknown. Here, we provide evidence that spontaneous fluctuations in alpha-band power and pupil size have distinct and separable effects on perception. From this, we postulate that measurements of pupil and alpha reflect different dimensions of internal state fluctuations (cortical excitement in alpha and arousal in pupil) which differentially shape trial-to-trial variability in our perception.

## 4. Materials and methods

### 4.1 Subjects

33 participants (24 female; mean age: 21.6) were recruited for this study from the University of California, Santa Cruz. All participants provided written consent and were compensated a $20 Amazon gift card as well as course credit for attending. All participants reported normal or corrected vision.

Three participants were excluded from the final analysis due to equipment malfunction (n=1), inability to sufficiently perform the task (n=1), and early termination of the study (n=1), leaving a sample size of 30 for the analysis reported here.

### 4.2 Stimuli

The experiment was implemented in MATLAB (The MathWorks Inc., 2022) and all stimuli were generated using Psychophysics Toolbox 3 (71). Stimuli were presented on a uniform gray background (approx. 50 cd/m^2^) on a gamma-corrected VIEWPixx EEG monitor (1920 x 1080 resolution, 120 Hz refresh rate). Participants were seated in a dimly lit room approximately 74 cm away from the screen with their head stabilized on a chin rest.

The target Gabor patch had a spatial frequency of 1.8 cycles per degree of visual angle (DVA) and spatial SD of 0.5 DVA. The target location was either in the upper right or upper left visual field, 5 DVA above the fixation and 5 DVA in either direction (see Figure 2).

An individual’s threshold level (i.e., the Michelson contrast value at which they reported perceiving the target about 50% of the time) was determined using a 1-up /1-down staircase procedure prior to the main task. The staircase relied on routines from the PALAMEDES toolbox (72). The threshold level was then multiplied by a range of values (0, 0.6, 0.8, 1, 1.4, 1.8) to produce a set of contrast levels spanning the CRF, including stimulus absent (0).

### 4.3 Procedure

Participants performed 800 trials of a yes-no visual detection task with confidence ratings. A target Gabor was presented for 8 ms at a random location in the visual field (upper left or upper right) and random contrast level (sampled with equal probability from the range specified above). After a 600ms delay, participants were cued via fixation color change to provide their detection and confidence response via a single button press. The right hand indicated that the participant perceived the target with four confidence levels (1 = “I’m guessing I saw the target”, 4 = “I’m certain I saw the target”) and the left hand indicated that they did not perceive the target with reciprocal confidence levels (see Figure 2).

Prior to the main task, participants performed one practice block (50 trials) of the task as well as one block (100 trials) of the up/down staircasing procedure. The average of the last 30 reversals provided the estimate of an individual’s threshold value (mean threshold value = 18.1% contrast). Total experiment time was approximately 3 hours.

### 4.4 EEG acquisition and preprocessing

EEG data was recorded using a 64 channel Ag/AgCl gel-based active electrode system (actiCHamp Plus, Brain Products GmbH, Gilching, Germany). The data was preprocessed using custom MATLAB scripts (version R2022a) and the EEGLAB toolbox (73). First, a high-pass filter was applied at 0.1 Hz, the data were down sampled to 500 Hz, and then re-referenced to the median of all electrodes. Then, the epoched data was visually inspected and trials with excessive noise, muscle artifacts, or ocular artifacts were rejected. Because we were specifically interested in spontaneous pupil activity and its link to EEG signals, we did not use ICA to remove eye-blink artifacts but instead rejected any trial with an eye blink or eye movement appearing within a one second window centered on stimulus onset. An average of 125 trials were removed. Channels with excessive noise were removed and spherically interpolated (average of 4 channels). Lastly, the data were re-referenced to the average.

### 4.5 Single trial pre-stimulus alpha power and residual alpha power

Single trial pre-stimulus alpha power was estimated using a fast Fourier transform (FFT) on a pre-stimulus window of -450ms to 50ms relative to stimulus onset (as close as we could estimate to stimulus processing but prior to any evoked responses). Single-trial pre-stimulus signals were Hamming-tapered and zero padded (by a factor of 5) to increase frequency resolution. The power spectrum was averaged over the six occipital electrodes showing highest 8-12 Hz power at the group level (see Figure 3B) as well as over each individual’s peak alpha frequency (IAF) +/- 2 Hz. For one participant who did not show a peak in the power spectrum in that range, an IAF of 10 was assigned.

Once alpha power was determined for each trial, these data were binned into four quartiles of ascending power (1 = lowest power trials, 4 = highest power trials). Primary analyses in this study considered the two most extreme bins, i.e., 1 and 4, as the high and low pre-stimulus alpha power conditions. Behavioral data were then sorted according to these high and low alpha bins.

Alpha power independent of pupil size was estimated by taking the single trial residuals of a linear model predicting pre-stimulus alpha power from pre-stimulus pupil size. The residuals of this model were then grouped in the same manner as alpha power (i.e., 4 bins of ascending power) for the purposes of examining behavior in these bins.

### 4.6 Pupillometry acquisition and preprocessing

Eye-tracking data was recorded using a binocular 2000 Hz TRACKPixx3 tabletop eye-tracker placed approximately 55 cm in front of the participant and below the stimulus presentation monitor. The eye-tracker was calibrated at the beginning of each block using the default TRACKPixx3 calibration routine. Triggers sent into the EEG system were synchronized with the eye data using a DATAPixx3. The epoched eye data were then aligned with the EEG data and any trials rejected in the EEG preprocessing stage were also removed from the eye data.

### 4.7 Single-trial pupil size

The average pre-stimulus pupil size on a given trial was determined by first assessing the median pupil value of both eyes between a -450 ms to 50 ms pre-stimulus window. We then checked that the median value was within a biologically plausible range (10 < value < 40 pixels), otherwise if both pupils did not fall within the range of plausible pupil sizes, that trial was omitted from the analysis (grand total of 70 omitted trials). If the value of both pupils fell within the plausible range, the average of both eyes was used to estimate the given trial’s pupil size. And if only one pupil was within the plausible range, then that one constituted the median pupil size of the given trial. The single trial average pupil sizes were then z-scored within each block, to account for possible block-wise differences in the efficacy of the eye-tracker calibration or the head position of the participants. The z-scored, single-trial pupil size estimate was then binned into four quartiles of ascending size (1 = smallest pupil size trials, 4 = largest pupil size trials).

### 4.8 Signal detection measurements

#### Detection reports

For each contrast level and alpha/pupil bin, we computed the proportion of trials where the subject reported detecting the stimulus. On stimulus absent trials, this proportion corresponds to the false alarm rate (FAR). For all other contrast levels this proportion corresponds to the hit rate (HR). For all SDT measures, a log-linear correction (74) was applied to the detection reports in order to accommodate perfect performance at any given contrast level and alpha bin.

#### Criterion

Criterion (c) was calculated by adding the z-transformed HR to the z-transformed FAR and multiplying it by -1/2. The formula for calculating c is as follows:

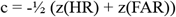

#### Sensitivity

Sensitivity (d’) was calculated by subtracting the z-transformed FAR by the z-transformed HR. The formula for calculating d’ is:

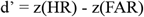

#### Generalized linear model (GLM)

The GLM for an analogous parameterization of SDT measurements uses a probit link function for estimating the probability of the response “seen” based on stimulus presence (stimpres), pre-stimulus alpha power (alphapow), and the interaction term of these two (stimpres:alphapow). The coefficients outputted by this model are used for the SDT calculations.

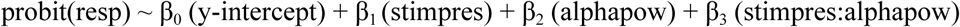

The intercept (β_0_) of this model can be understood in the context of whether or not the stimulus was presented (the effect of which is β_1_) and is represented as either 0 (absent) or 1 (presented). In the case of stimulus absence, β_0_ represents the FAR. In the case of stimulus presence, β_0_+ β_1_ represents the HR. The effect that alpha power has on the probability of saying “seen” – irrespectively of stimulus presence – is represented by β_2_. And the effect that alpha power has on reports given the stimulus presence is β_3_.

#### GLM criterion

C was computed by taking the negative of the sum of the intercept and alpha power coefficient and subtracting that by half the product of the stimulus presence coefficient by the interaction coefficient.

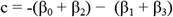

#### GLM sensitivity

d’ was calculated by adding the stimulus presence coefficient to the interaction coefficient.

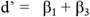

#### GLM alpha power on criterion

The effect of alpha power on criterion was computed by subtracting the negative of the alpha power coefficient from half of the interaction coefficient.

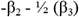

#### GLM alpha power on sensitivity

The effect of alpha power on sensitivity was the interaction coefficient, β_3._

## Acknowledgements

We sincerely thank Nitu Gupta, Antonia Gergen, Alex Feghhi, and Shirin Afrakhteh for help with data collection.

